# Unveiling targeted cell-free DNA methylation regions through paired methylome analysis of tumor and normal tissues

**DOI:** 10.1101/2023.06.27.546654

**Authors:** Tingyi Li, Krupal B Patel, Xiaoqing Yu, Sijie Yao, Liang Wang, Christine H Chung, Xuefeng Wang

## Abstract

Liquid biopsy analysis of cell-free DNA (cfDNA) has revolutionized cancer research by enabling non-invasive assessment of tumor-derived genetic and epigenetic changes. In this study, we conducted a comprehensive paired-sample differential methylation analysis (psDMR) on reprocessed methylation data from two large datasets, CPTAC and TCGA, to identify and validate differentially methylated regions (DMRs) as potential cfDNA biomarkers for head and neck squamous cell carcinoma (HNSC). Our hypothesis is that the paired sample test provides a more suitable and powerful approach for the analysis of heterogeneous cancers like HNSC. The psDMR analysis revealed a significant number of overlapped hypermethylated DMRs between two datasets, indicating the reliability and relevance of these regions for cfDNA methylation biomarker discovery. We identified several candidate genes, including *CALCA, ALX4*, and *HOXD9*, which have been previously established as liquid biopsy methylation biomarkers in various cancer types. Furthermore, we demonstrated the efficacy of targeted region analysis using cfDNA methylation data from oral cavity squamous cell carcinoma and nasopharyngeal carcinoma patients, further validating the utility of psDMR analysis in prioritizing cfDNA methylation biomarkers. Overall, our study contributes to the development of cfDNA-based approaches for early cancer detection and monitoring, expanding our understanding of the epigenetic landscape of HNSC, and providing valuable insights for liquid biopsy biomarker discovery not only in HNSC and other cancer types.

## INTRODUCTION

In recent years, the field of cancer research has witnessed a promising advancement with the emergence of cell-free DNA (cfDNA) analysis, which has opened new avenues for non-invasive assessment of tumor-derived genetic and epigenetic alterations. Liquid biopsies from cancer patients, such as plasma, urine, and saliva, serve as reservoirs for cell-free DNA (cfDNA) that carry fragments of circulating tumor DNA (ctDNA), which are shed into circulation because of cancer cell death and secretion. cfDNA found in health or cured individuals predominantly originates from hematopoietic cells, specifically lymphocytes and erythrocyte progenitors. Therefore, the cfDNA obtained from individuals belonging to different clinical groups, including those with varying tumor burden, stage, or treatment regimens, as well as patients throughout their treatment journey, may exhibit considerable variations in both concentration and composition. In general, cfDNA methylation exhibits greater sensitivity in detecting early-stage cancer due to extensive epigenetic alternations occurring during in the initial phases of cancer progression, as opposed to mutation-based methods^1^. By leveraging the distinct methylation signatures found in cfDNA, clinicians and researchers can gain valuable insights into tumor progression, response to treatment, and even the emergence of treatment-resistant clones. Moreover, the non-invasive nature of cfDNA analysis allows for repeated testing and real-time monitoring of disease dynamics, offering a minimally invasive approach that circumvents the limitations and risks associated with traditional tissue biopsies. These advancements thus paved the way for personalized medicine, where tailored treatment strategies can be developed based on the unique cfDNA methylation profiles of individual patients.

There are many commonly accepted cfDNA methylation biomarkers that have been identified and validated across various types of cancer. For example, the hypermethylation of gene *SEPTIN9* (*SEPT9*) has been extensively studied as an early screening biomarker for colorectal cancer^2^; methylation status of *SHOX2* has been utilized in both lung cancer^3,4^ and colon cancer^5^ detection; *MGMT* has been studied in relation to brain tumors^6^; *RASSF1A* has shown as a potential candidate for pan-cancer biomarker^7,8^, aiding both cancer diagnosis and prognosis^9^. Emerging evidence also indicates that the methylation patterns of long interspersed nuclear element (LINE-1) hold promise as a new global cfDNA biomarker for cancer hallmark in multiple cancers^10,11^. Considering the intrinsic heterogeneity of cancer and the tissue-specific nature of DNA methylation, it is anticipated that the inclusion of multiple biomarkers in a predictive model will enhance both the sensitivity and specificity of cancer detection. Therefore, a panel of cfDNA methylation biomarkers should be tailored for each cancer type or even subtype, ideally also taking into account the technical biases and preferences associated with the specific cfDNA methylation profiling technologies being utilized.

Despite the capability of modern genome-wide cfDNA methylation assays to provide a holistic assessment of sites in a single experiment, it is important to note that many successful studies in cfDNA continue to rely heavily on candidate biomarkers derived from the tumor tissue data. In our opinion, this is attributed to three main reasons: Firstly, the relatively sample size and the temporal heterogeneity in current cfDNA studies poses limitations on effective training; Secondly, the differential methylation analysis by comparing methylome of tumor and normal tissues remains a powerful approach for identifying cancer-specific biomarkers; Thirdly, given the low amount of cfDNA in liquid biopsies, the selective enrichment of cfDNA fragments over the background non-cancer-specific regions will greatly increase the detection sensitivity in most techniques. Our literature review indicates the methylation data generated from The Cancer Genome Atlas (TCGA) project remain the mainstay for the initial methylation region discovery in most cancers. For example, in a study for discovering non-invasive blood-based markers for lung cancer^12^, a panel of 12,899 pre-selected cancer-specific informative regions was first constructed based on TCGA lung cancer database.

Due to the initial emphasis of the TCGA project on investigating the landscape of cancer genomes, the methylation data in TCGA has predominantly been obtained from tumor tissues. For example, within the TCGA Head-Neck Squamous Cell Carcinoma (TCGA-HNSC) dataset, there are 528 tumor tissues available for methylation data, whereas only 50 normal tissue samples were profiled with methylation. In our previous study^13^, we discovered that the more significant differential methylated regions (DMRs) can be detected by including only those samples with paired tumor and normal tissues in the analysis. This finding underscores the critical importance of conducting paired-sample differential methylation analysis (psDMR), as it enhances the statistical power and reliability of identifying candidate regions. In the context of cancers such as HNSC, which exhibit remarkable sample heterogeneity due to tumor site and molecular subtypes, the self-controlled psDMR becomes even more crucial. By comparing methylation patterns within paired samples, we can effectively account for the inherent variability and confounding factors associated with such heterogeneity, leading to more accurate and insights into the epigenetic dynamics of HNSC.

In this study, our main objective is to conduct a thorough psDMR analysis of the methylation data in HNSC with the aim of developing an enhanced resource in the form of methylation panel that can be effectively utilized in targeted cfDNA methylation studies. To achieve this, we embark on a systematic approach. We first revisit the TCGA data and conduct psDMR analysis based on the updated HumanMethylation450 (450K) methylation data sourced from HNSC-TCGA, which has been reprocessed using the SeSAMe pipeline^14^, ensuring refined data quality. Furthermore, we expand our analysis by incorporating the MethylationEPIC (EPIC) methylation data obtained from matched tumor-normal samples in the CPTAC-HNSC study^15^. By conducting a similar psDMR analysis on this independent dataset, we aim to validate our findings from the original TCGA data and identify additional methylation sites of interest. Lastly, we demonstrate the application of the targeted psDMR regions based on two cfDNA studies. These studies provide real-world examples of how the identified psDMR regions can be effectively employed in investigating cfDNA methylation patterns for cancer detection, especially in scenarios with limited sample sizes. Importantly, the comprehensive biomarker prioritization strategy and findings presented in this study have the potential to extend beyond HNSC, as they can be applied to the analysis of methylation data in other cancer types. Collectively, by demonstrating the practical utility and effectiveness of the targeted psDMR analysis, we contribute to the advancement of cfDNA-based approaches for early cancer detection and monitoring, broadening the scope of their impact in the field of cancer research.

## METHOD AND RESULTS

### psDMR on CPTAC-HNSC reveals candidate regions for cfDNA biomarkers

The psDMR analysis conducted on the methylation data obtained from 42 paired tumor and normal tissues in CPTAC-HNSC cohort successfully identified a total of 769 significantly differentially methylated regions with FWER <0.05 (using the modified bumphunter algorithm implemented in *ChAMP)*. Out of these 769 regions, 573 of them were found to be located in promoter regions, corresponding to 561 unique genes. Within this gene set, it was observed that 11 genes, such as *PAX6, WT1*, and *CA10*, harbored multiple significant DMRs. The detailed DMR information regarding all significant DMRs are listed in **Supplementary Table 1**, while the top 30 DMRs and their associated names are highlighted in **Table 1**. Notably, among the top 30 DMRs, as well as among all significantly differentially methylated regions, a significant majority displayed hypermethylation when comparing tumor vs. normal tissues. Specifically, 83.3% (n=25) of the regions in the top 30 DMRs and 73.1% (n=562) of all significant DMRs exhibited hypermethylation.

**Table 1.** Top 30 genomic regions identified using the paired DMR analysis on CPTAC-HNSC data.

| DMR (CPTAC) | Chr | Start | End | Gene symbol<sup>†</sup> | EntrezID | No. of CpGs | FWER | FWER.area | DMR state |
| --- | --- | --- | --- | --- | --- | --- | --- | --- | --- |
| 1 | chr11 | 2321770 | 2323459 | *C11orf21* | 29125 | 28 | <0.001 | 0.032 | *Hypo* |
| 2 | chr11 | 14993378 | 14995770 | *CALCA*<sup>†</sup> | 796 | 40 | <0.001 | 0.032 | Hyper |
| 3 | chr11 | 44332385 | 44333192 | *ALX4*<sup>†</sup> | 60529 | 28 | <0.001 | 0.04 | Hyper |
| 4 | chr10 | 60936025 | 60937501 | *PHYHIPL* | 84457 | 20 | <0.001 | 0.044 | Hyper |
| 5 | chr12 | 22093960 | 22095330 | *ABCC9* | 10060 | 16 | <0.001 | 0.044 | Hyper |
| 6 | chr2 | 198650603 | 198651590 | *BOLL*<sup>†</sup> | 66037 | 16 | <0.001 | 0.044 | Hyper |
| 7 | chr3 | 147125712 | 147127193 | *ZIC1*<sup>†</sup> | 7545 | 23 | <0.001 | 0.044 | Hyper |
| 8 | chr5 | 115296978 | 115299088 | *LVRN* | 206338 | 19 | <0.001 | 0.048 | Hyper |
| 9 | chr10 | 118031632 | 118034031 | *GFRA1* | 2674 | 27 | <0.001 | 0.052 | Hyper |
| 10 | chr16 | 51183988 | 51185672 | *SALL1* | 6299 | 25 | <0.001 | 0.052 | Hyper |
| 11 | chr7 | 27281216 | 27282987 | *EVX1* | 2128 | 27 | <0.001 | 0.052 | Hyper |
| 12 | chr18 | 904851 | 905611 | *ADCYAP1*<sup>†</sup> | 116 | 15 | <0.001 | 0.056 | Hyper |
| 13 | chr19 | 52390810 | 52391789 | *ZNF577*<sup>†</sup> | 84765 | 13 | <0.001 | 0.056 | Hyper |
| 14 | chr8 | 23563572 | 23565174 | *NKX2-6*<sup>†</sup> | 137814 | 16 | <0.001 | 0.056 | Hyper |
| 15 | chr12 | 95941571 | 95943131 | *USP44*<sup>†</sup> | 84101 | 18 | <0.001 | 0.056 | Hyper |
| 16 | chr14 | 101489415 | 101490253 | *MIR411* | 693121 | 17 | <0.001 | 0.056 | *Hypo* |
| 17 | chr5 | 140810051 | 140811520 | *PCDHGA12*<sup>†</sup> | 26025 | 17 | <0.001 | 0.06 | Hyper |
| 18 | chr2 | 176980673 | 176981919 | *HOXD10*<sup>†</sup> | 3236 | 12 | <0.001 | 0.064 | Hyper |
| 19 | chr6 | 133562196 | 133562776 | *EYA4*<sup>†</sup> | 2070 | 19 | <0.001 | 0.068 | Hyper |
| 20 | chr6 | 33047944 | 33048879 | *HLA-DPB1* | 3115 | 17 | <0.001 | 0.076 | Hyper |
| 21 | chr14 | 36982447 | 36983791 | *SFTA3* | 253970 | 15 | <0.001 | 0.076 | Hyper |
| 22 | chr19 | 38182773 | 38183460 | *ZNF781*<sup>†</sup> | 163115 | 12 | <0.001 | 0.076 | Hyper |
| 23 | chr17 | 2699249 | 2700093 | *RAP1GAP2* | 23108 | 12 | <0.001 | 0.076 | *Hypo* |
| 24 | chr5 | 113697330 | 113698847 | *KCNN2* | 3781 | 17 | <0.001 | 0.076 | Hyper |
| 25 | chr2 | 176986535 | 176987605 | *HOXD9*<sup>†</sup> | 3235 | 12 | <0.001 | 0.076 | Hyper |
| 26 | chr19 | 58446312 | 58446988 | *ZNF418* | 147686 | 11 | <0.001 | 0.076 | Hyper |
| 27 | chr14 | 101531403 | 101532352 | *MIR409* | 574413 | 18 | <0.001 | 0.076 | *Hypo* |
| 28 | chr17 | 5402883 | 5404581 | *LOC728392* | 728392 | 16 | <0.001 | 0.076 | Hyper |
| 29 | chr19 | 58220080 | 58220955 | *ZNF154*<sup>†</sup> | 7710 | 11 | <0.001 | 0.076 | Hyper |
| 30 | chr14 | 101491151 | 101492406 | *MIR380* | 494329 | 14 | <0.001 | 0.076 | *Hypo* |

<sup>†</sup>Genes that have been previously established as cfDNA or liquid biopsy methylation biomarkers in at least one cancer type.

In the top 30 DMRs, we observed that 14 of them are located within genes that have been previously established as liquid biopsy methylation biomarkers in at least one cancer type to date, indicating the relevance and importance of these genes as reliable cancer-specific markers. These genes include *CALCA, ALX4, BOLL, ZIC1, ADCYAP1, ZNF577, NKX2-6, USP44, PCDHGA12,HOXD10, EYA4, ZNF781, HOXD9*, and *ZNF154*. The gene *CALCA*, encoding the calcitonin gene-related peptide that possess tumor suppressive potential, has been documented as being hypermethylated in several malignancies including head and neck cancer^16^. *CALCA* has also been reported as an effective cfDNA methylation biomarkers for early diagnosis of thyroid cancer^17^, ovarian cancer^18^, and testicular germ cell tumor^19^. The genes *ALX4* and *ZIC1* are transcription factors that play pivotal roles in regulation of embryonic development, and they are identified as epigenetically downregulated tumor suppressor^20,21^. The genes *HOXD9* and *HOXD10* belong to HOX gene cluster, and the methylation status of this gene family have been extensively explored as liquid biomarker in multiple cancers^22^. Notably, in our previously study we also reported the diagnostic potential of these two genes using plasma samples from oral cavity squamous cell carcinoma (OCSS) ^13^. Other HOX genes identified from the significant DMRs include *HOXA2, HOXA3, HOXA7, HOXA9, HOXB1, HOXC4, HOXC6, HOXC9, HOXD4*, and *HOXD12*. Another noteworthy discovery that strongly aligns with previous analysis in OCSS is the identification of ZNF family genes (e.g., *ZNF577, ZNF781, ZNF418 and ZNF154*) in the top-ranked list. The entire significant DMR list encompasses a total of 46 ZNF family genes. These genes are recognized for their roles in transcriptional regulation and tumor suppressor activity and have gained increased attention in the context of head and neck cancers^13,23^.

An intriguing observation within the top 30 DMR list is the presence of three hypomethylated region (out of five) derived from microRNA genes, specifically *MIR411, MIR409* and *MIR380*. Furthermore, a total of 21 significant DMRs were identified in microRNAs. Among them, additional microRNAs with hypomethylation status include *MIR381, MIR376C, MIR668, MIR127, MIR495, MIR124-3, MIR10B, MIR382*, and *MIR496*.

### Consistent DMRs validated by TCGA-HNSC psDMR analysis

We performed the similar analysis of the TCGA data by conducting psDMR analysis on the 450K array methylation data obtained from the paired samples in HNSC-TCGA. The beta-values (downloaded from GDC Data Portal) were processed via the same SeSAMe pipeline utilized in CPTAC, thus minimizing data inconsistency arising from different bioinformatics pipelines. The psDMR analysis on the TCGA data identified a total of 521 significantly differentially methylated regions with FWER <0.05. Among them, 359 regions found to be located within gene promoter regions, corresponding to 355 unique genes. The top 30 DMRs from the TCGA analysis and their associated names are listed in **Table 2**. Out of those DMRs, 29 regions (excluding *C11orf21*) and a notable majority of all significant DMRs, specifically 90.3% (324/359), displayed hypermethylation. The overall results demonstrated a high level of consistency between the findings from the CPTAC and TCGA. As indicatecd in the gene symbol column of the table, 28 out of top 30 DMRs in TCGA were also found significant in CPTAC. Even more strikingly, among these, a total of 12 DMR genes were consistently identified as top 30 DMRs in both the TCGA and CPTAC analyses. These genes are *ALX4, CALCA, ADCYAP1, PCDHGA12, C11orf21, ZIC1, ABCC9, EVX1, GFRA1, PHYHIPL, HOXD9*, and *NKX2-6*. Additionally, another 10 DMR genes in the were identified as top 100 DMRs in the CPTAC analysis, namely *EDNRB, CCNA1, ZIM2, MARCHF11, PENK, MIR124-2, ITGA8, ASB3, ZNF542P* and *ZNF454*. By specifically examining DMRs located in promoter regions, we identified a notable overlap of 49 overlapped genes among the top 100 DMRs from two datasets and a total of 264 overlapped genes from the lists of all significant DMRs. This remarkable overlap highlights the robustness and reproducibility of psDMR results across independent datasets, providing further support for their potential translational significance in the context of HNSC liquid biopsy studies. Furthermore, the analysis revealed a significant presence of HOX and ZNF family genes in both TCGA and CPTAC datasets. Alongside HOXD9, which appeared in the top 30 list from the TCGA analysis, several other HOX genes including *HOXD10, HOXD12, HOXA7, HOXA9, HOXB1, HOXC4* and *HOXC6* were also identified. Apart from *ZNF542P* and *ZNF454*, an additional set 34 ZNF genes were found to harbor significant DMRs in the TCGA analysis. A total of 11 microRNAs were identified in the significant DMR analysis with TCGA data. Five of them displayed hypomethylation status; and four out of these five, namely *MIR411, MIR495, MIR496*, and *MIR668*, were also identified in CPTAC data.

**Table 2.** Top 30 genomic regions identified using the paired DMR analysis on TCGA-HNSC data.

| DMR (TCGA) | Chr | Start | End | Gene Symbol* | EntrezID | No. of CpGs | FWER | FWER. area | DMR state |
| --- | --- | --- | --- | --- | --- | --- | --- | --- | --- |
| 1 | chr11 | 44332332 | 44333192 | **ALX4** *** | 60529 | 30 | <0.001 | 0.004 | Hyper |
| 2 | chr11 | 14993378 | 14995770 | **CALCA** *** | 796 | 43 | <0.001 | 0.004 | Hyper |
| 3 | chr7 | 27224954 | 27226148 | **HOXA11-AS** | 221883 | 25 | <0.001 | 0.004 | Hyper |
| 4 | chr13 | 78492678 | 78493712 | **EDNRB** ** | 1910 | 32 | <0.001 | 0.004 | Hyper |
| 5 | chr15 | 26108084 | 26108948 | **ATP10A** | 57194 | 18 | <0.001 | 0.008 | Hyper |
| 6 | chr13 | 37004536 | 37005582 | **CCNA1** ** | 8900 | 24 | <0.001 | 0.008 | Hyper |
| 7 | chr18 | 904851 | 905611 | **ADCYAP1** *** | 116 | 14 | <0.001 | 0.008 | Hyper |
| 8 | chr5 | 140810051 | 140811642 | **PCDHGA12** *** | 26025 | 17 | <0.001 | 0.008 | Hyper |
| 9 | chr11 | 2321770 | 2323459 | **C11orf21** *** | 29125 | 30 | <0.001 | 0.008 | Hypo |
| 10 | chr3 | 147125712 | 147127193 | **ZIC1** *** | 7545 | 26 | <0.001 | 0.012 | Hyper |
| 11 | chr12 | 22093960 | 22095330 | **ABCC9** *** | 10060 | 13 | <0.001 | 0.012 | Hyper |
| 12 | chr4 | 4859772 | 4860655 | **MSX1** * | 4487 | 19 | <0.001 | 0.012 | Hyper |
| 13 | chr19 | 57351322 | 57352807 | **ZIM2** ** | 23619 | 23 | <0.001 | 0.012 | Hyper |
| 14 | chr5 | 16179633 | 16180419 | **MARCHF11** ** | 441061 | 13 | <0.001 | 0.012 | Hyper |
| 15 | chr7 | 27280914 | 27282444 | **EVX1** *** | 2128 | 22 | <0.001 | 0.012 | Hyper |
| 16 | chr10 | 118031632 | 118034031 | **GFRA1** *** | 2674 | 25 | <0.001 | 0.012 | Hyper |
| 17 | chr8 | 57358130 | 57359414 | **PENK** ** | 5179 | 13 | <0.001 | 0.012 | Hyper |
| 18 | chr10 | 60935663 | 60937257 | **PHYHIPL** *** | 84457 | 17 | <0.001 | 0.012 | Hyper |
| 19 | chr7 | 19157710 | 19158954 | **TWIST1** * | 7291 | 21 | <0.001 | 0.012 | Hyper |
| 20 | chr8 | 65291287 | 65292498 | **MIR124-2** ** | 406908 | 14 | <0.001 | 0.012 | Hyper |
| 21 | chr10 | 15760947 | 15762312 | **ITGA8** ** | 8516 | 13 | <0.001 | 0.012 | Hyper |
| 22 | chr2 | 54086854 | 54087552 | **ASB3** ** | 51130 | 13 | <0.001 | 0.016 | Hyper |
| 23 | chr13 | 108518955 | 108521062 | **FAM155A** * | 728215 | 16 | <0.001 | 0.02 | Hyper |
| 24 | chr7 | 37488162 | 37489005 | **ELMO1** * | 9844 | 15 | <0.001 | 0.024 | Hyper |
| 25 | chr2 | 176986460 | 176987605 | **HOXD9** *** | 3235 | 12 | <0.001 | 0.024 | Hyper |
| 26 | chr19 | 56879369 | 56879994 | **ZNF542P** ** | 147947 | 11 | <0.001 | 0.024 | Hyper |
| 27 | chr5 | 178367621 | 178368620 | **ZNF454** ** | 285676 | 11 | <0.001 | 0.024 | Hyper |
| 28 | chr8 | 23563572 | 23564717 | **NKX2-6** *** | 137814 | 12 | <0.001 | 0.024 | Hyper |
| 29 | chr7 | 15725086 | 15726821 | **MEOX2** * | 4223 | 15 | <0.001 | 0.024 | Hyper |
| 30 | chr19 | 56904580 | 56905152 | **ZNF582** * | 147948 | 11 | <0.001 | 0.024 | Hyper |

\*Genes validated by CPTAC dataset are highlighted: (\*\*\*) indicates top 30 DMR; (\*\*) indicates top 100 DMR; (\*) indicates significant DMR in the CPTAC-HNSC paired sample analysis.

As summarized in **Figure 1**, the comparative results of our study not only demonstrate the reliability of psDMR analysis across datasets but also contribute to expanding our understanding of the epigenetic landscape specific to HNSC.

**Figure 1.**
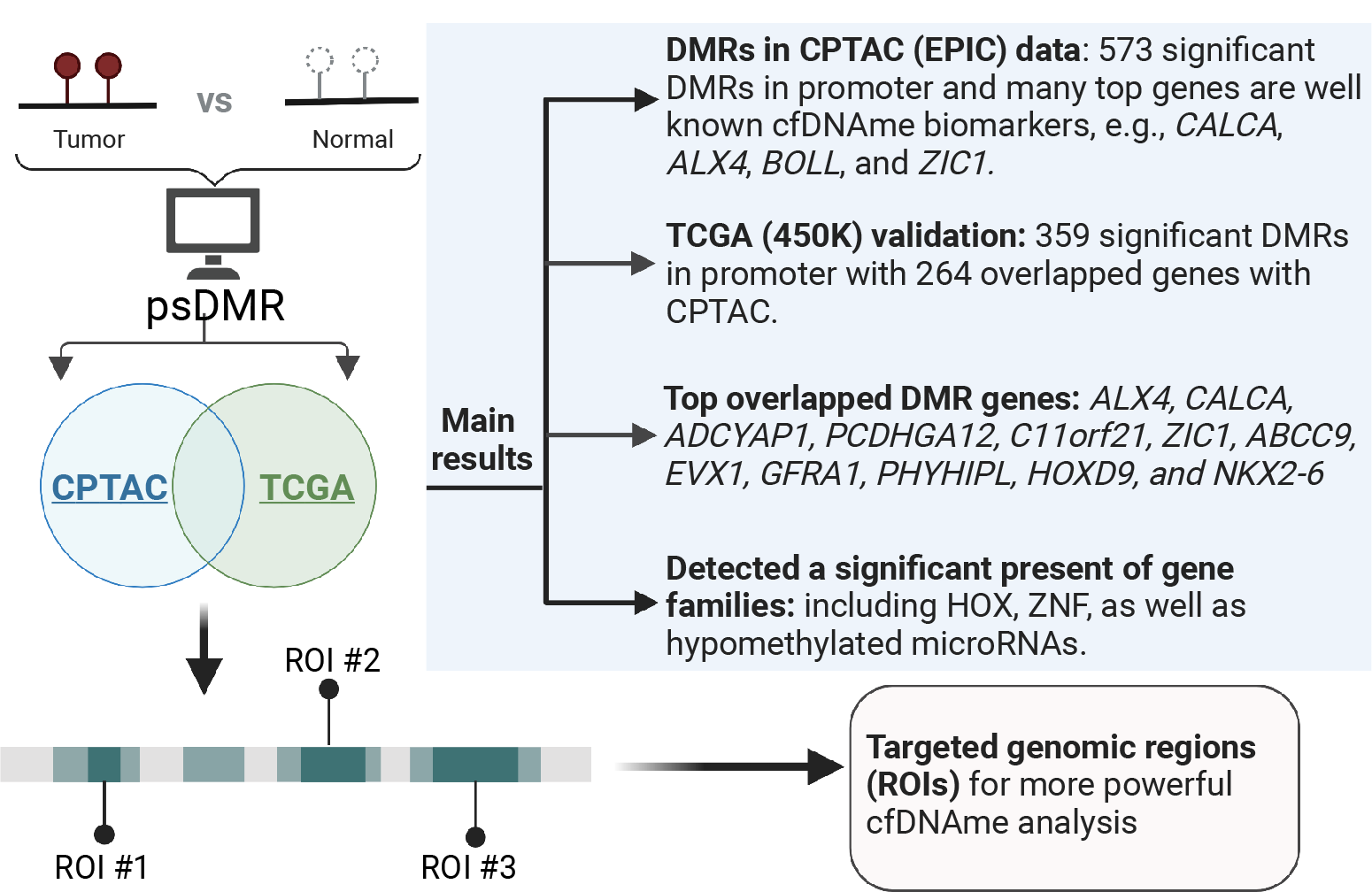
Workflow of paired sample DMR analysis and key findings in HNSC samples.

### Leveraging targeted regions in genome-wide cfDNA DMR analysis

In order to showcase the effectiveness of targeted regions in providing insights for liquid biopsy analysis in HNSC, we conducted a candidate region-based DMR analysis based on plasma cfMBD-seq obtained from eight patients diagnosed with OCSCC^13^. We perform genome-wide differential coverage analysis on cfDNA methylation data by only considering those regions are determined to be significant in the comparison of matched tumor-normal tissues from the CTPAC analysis. The overall analysis steps of paired sample DMR on cfDNA methylation data followed a similar approach to our previous analysis. Specifically, we employed “MEDIPS” to calculate normalized methylation levels in all targeted ROI (region of interest) regions. Subsequently, fitNBglm function from the “qsea” package was utilized to perform differential analysis for each genomic window. Due to the restricted sample size and the validation-focused nature of the study, we employed unadjusted p-values as a criterion for prioritizing the top differentially methylated regions (DMRs). Out of the analyzed regions, a total of 185 regions demonstrated statistical significance (*P*<0.05). The top five DMR regions from the cfDNA data are identified within the promoter regions of the following genes: *PENK, TRH, PIEZO2, CA8*, and *HPSE2*. Remarkably, among the top 30 DMR genes identified based on the CPTAC psDMR analysis, we observed that six of them, namely *ABCC9, NKX2-6, USP44, HOXD9, ZNF418* and *ZNF154*, are also present in the significant DMR list obtained from the cfDNA targeted DMR analysis. The consistency observed in the overall pattern across patients, as depicted in **Figure 2A**’s waterfall plot, further reinforces the encouraging nature of these results in alignment with our previous analysis.

**Figure 2.**
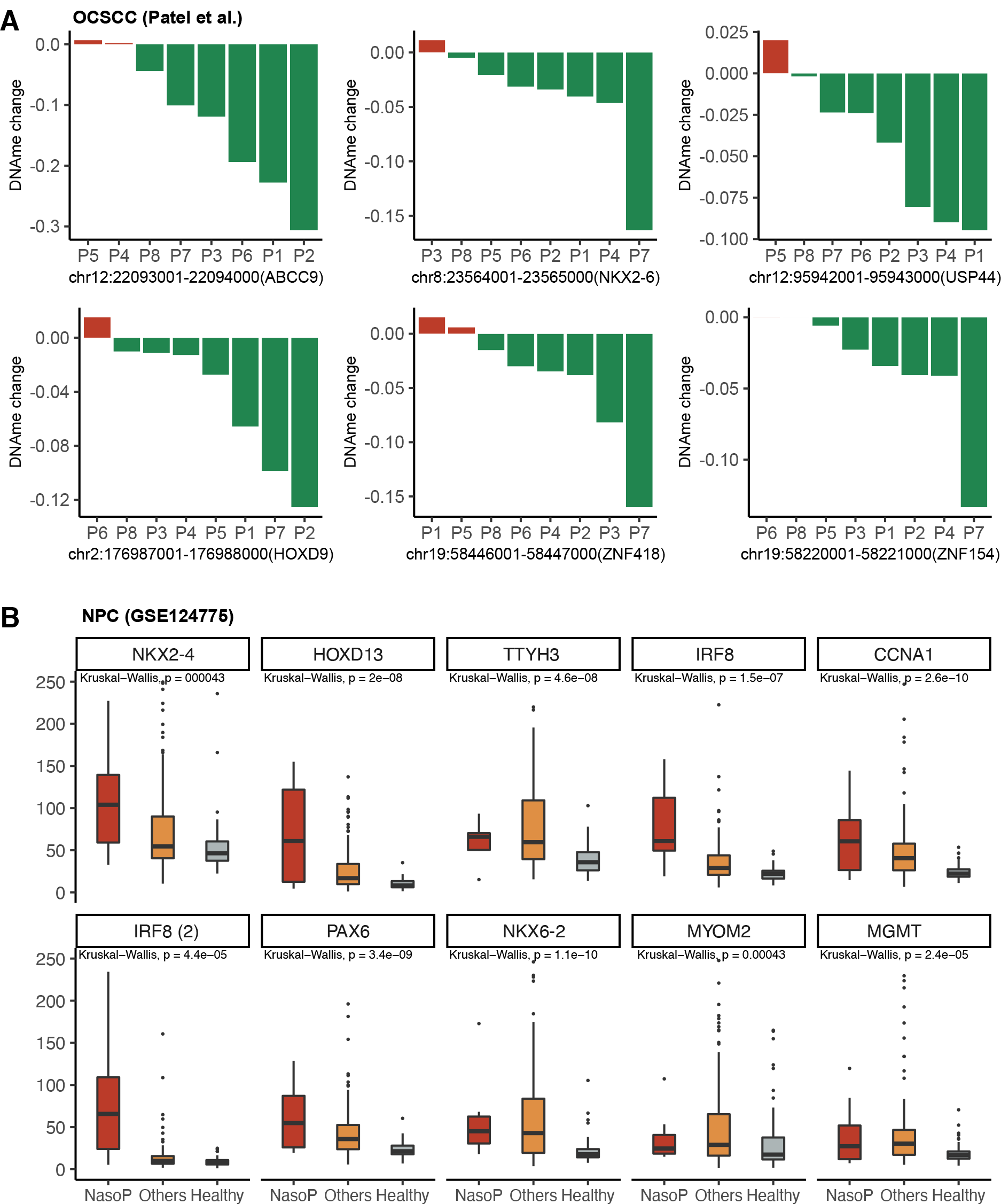
Validation of the top DMRs based on plasma cfDNA methylation data obtained from two HNSC studies: (A) OSCC and (B) NPC.

In another plasma-based cfDNA methylation from multiple cancer types including nasopharyngeal carcinoma (GSE124775), we found 17 genes (out of 91 targeted genes available in the dataset) overlapped with DMR genes identified based on the CPTAC psDMR analysis. The shared genes are *CCNA1, NKX2-4, FOXG1, PAX6, CNR1, NKX6-2, TWIST1, MYOM2, NKX6-2, IRF8, RUNX3, FOXG1, HOXD13, MGMT, TWIST1, TTYH3*, and *PRDM2*. To facilitate the cancer-specific comparison, we divided all the samples into three main groups: nasopharyngeal, other malignancies and healthy. We further excluded genes that showed low values across three main groups, resulting in the identification of 9 genes (2 regions from *IRF8*). The methylation levels of these genes are depicted in **Figure 2B**. All these genes showed significantly elevated methylation levels in cancer samples versus healthy samples. More importantly, many of them (e.g. *HOXD13, IRF8*, and *CCNA1*), also demonstrated notably higher methylation specifically in NPC samples when compared to other malignancies. This finding strongly supports the notion that these candidate genes hold promise as potential biomarkers specific to HNSC.

## CONCLUSIONS

In conclusion, our study utilizing psDMR analysis on the CPTAC-HNSC and TCGA-HNSC datasets successfully identified reliable differentially methylated regions, which could inform more powerful cfDNA methylation biomarker discovery. Within the top DMRs, a significant majority displayed hypermethylation in tumor tissues compared to normal tissues, indicating that the targeted regions identified based on the methylation array (EPIC and 450K) will be relevant to enrichment-based cfDNA methylation methods, which target hypermethylation only. Moreover, many top DMRs were found located in genes that have been previously established as liquid biopsy methylation biomarkers in various cancer types, underscoring their reproducibility and importance as reliable cancer-specific markers. Consistency was also observed between the CPTAC and TCGA in terms of the HOX and ZNF gene families, as well as microRNAs, emphasizing their involvements in the epigenetic landscape of HNSC. Furthermore, we demonstrated the efficacy of targeted region analysis using cfDNA methylation data from OCSCC patients. The analysis identified several significant DMRs in regions previously determined to be significant in the CPTAC analysis. Consistency was also observed with another cfDNA study encompassing multiple cancer types, including NPC. We identified 9 overlapping gens with the DMR genes from the CPTAC analysis. Notably, the methylation levels of many genes not only showed significant elevation in cancer samples compared to healthy samples, but also showed notably higher levels in NPC samples compared to other cancer types, supporting their potential as HNSC-specific biomarkers. Overall, our comprehensive analysis demonstrates the reliability and translational significance of psDMR analysis in HNSC, providing insights into the epigenetic landscape and potential liquid biopsy biomarkers for this cancer type.

